# Metagenomic analysis of individual mosquitos reveals the ecology of insect viruses

**DOI:** 10.1101/2023.08.28.555221

**Authors:** Yuan-fei Pan, Hailong Zhao, Qin-yu Gou, Pei-bo Shi, Jun-hua Tian, Yun Feng, Kun Li, Wei-hong Yang, De Wu, Guangpeng Tang, Bing Zhang, Zirui Ren, Shiqin Peng, Geng-yan Luo, Shi-jia Le, Gen-yang Xin, Jing Wang, Xin Hou, Min-wu Peng, Jian-bin Kong, Xin-xin Chen, Chun-hui Yang, Shi-qiang Mei, Yu-qi Liao, Jing-xia Cheng, Juan Wang, Chaolemen, Yu-hui Wu, Jian-bo Wang, Tongqing An, Xinyi Huang, John-Sebastian Eden, Jun Li, Deyin Guo, Guodong Liang, Xin Jin, Edward C. Holmes, Bo Li, Daxi Wang, Junhua Li, Wei-chen Wu, Mang Shi

## Abstract

Mosquito transmitted viruses are responsible for an increasing burden of human disease. Despite this, little is known about the diversity and ecology of viruses within individual mosquito hosts. Using a meta-transcriptomic approach, we analysed the virome of 2,438 individual mosquitos (79 species), spanning ∼4000 km along latitudes and longitudes in China. From these data we identified 393 core viral species associated with mosquitos, including seven (putative) arbovirus species. We identified potential species and geographic hotspots of viral richness and arbovirus occurrence, and demonstrated that host phylogeny had a strong impact on the composition of individual mosquito viromes. Our data revealed a large number of viruses shared among mosquito species or genera, expanding our knowledge of host specificity of insect-associated viruses. We also detected multiple virus species that were widespread throughout the country, possibly facilitated by long-distance mosquito migrations. Together, our results greatly expand the known mosquito virome, linked the viral diversity at the scale of individual insects to that at a country-wide scale, and offered unique insights into the ecology of viruses of insect vectors.

## INTRODUCTION

Mosquitos (Diptera: Culicidae) are vectors for various arthropod-borne viruses that infect humans and other animals, including dengue virus, Chikungunya virus, and Zika virus^1^. In addition to their role in disease transmission, mosquitos also harbour a highly diverse virome, encompassing many “insect-specific” viruses that are not associated with the infection of vertebrates^2,3^. Although these insect-specific viruses do not directly impact public health, some are known to influence the transmission of arboviruses^4^. Despite this, we lack key epidemiological and ecological information on the viruses associated with insect vectors in general, including their distribution, prevalence, co-infection, transmission, and host specificity.

Metagenomic studies of individual insects are necessary to reveal the epidemiology and ecology of viruses without *a priori* knowledge of which viruses may be present^5^. Individual animal virome data sets are also valuable for investigating both virus-virus and virus-host interactions^4,6^. Many metagenomics studies have pooled individual insects by species or sampling location^7–10^. Although pooling is an efficient means to explore viral diversity, it inevitably hinders mechanistic insights into the viral ecology and evolution. Only five studies to date have characterized the virome of individual insects^11–15^. While these studies have provided interesting insights into viral prevalence, coinfection, and the drivers of virome compositions, it remains uncertain whether the patterns and drivers revealed are generalizable due to their small sample size and limited focus on specific areas and species. Hence there is still an important need for virome data sets from individual insect vectors.

It is also important to establish links between viral diversity at the scale of individual animals and at larger scales, such as an entire country^16–18^. By including insect individuals from different regions, a more generalizable perspective of viral ecology may be obtained. In addition, collecting individual insects in an unbiased manner from diverse climatic zones and habitats and comparing their virome compositions is crucial for understanding the biogeography of insect-associated viruses. Comparing the diversity and prevalence of viruses among areas, especially arboviral species, could identify potential hotspots for vector-borne diseases emergence^19,20^. This information could then guide disease surveillance. Such data is currently lacking^21^.

The present study addresses these substantive gaps by characterizing the virome of 2,438 individual mosquitos from diverse habitats across China. We aimed to reveal the patterns and drivers of viral diversity at both the finest (i.e., individual) scale, and more broadly across China, and determine the ecological properties of mosquito-associated viruses. Specifically, we examined the potential environmental (climate and land-use) and host (mosquito species) drivers of virus diversity and identified potential diversity hotspots. In addition, we investigated the effect of host phylogeny and spatial distance on virus transmission, assessing the host specificity of mosquito-associated viruses. Finally, we explored the biogeographic patterns (distribution of viruses throughout the entire country) of mosquito-associated viruses and investigated the role of mosquito movement in shaping virus distribution.

## RESULTS

### Characterisation of Individual Mosquito Viromes

We conducted meta-transcriptomic (i.e., total RNA) sequencing of 2,438 individual mosquitos collected from diverse habitats across China between 2018 and 2021 (Fig. 1; Supplemental Data 1). This generated 9.8 billion non-rRNA reads (3.6 million per sample on average), which were *de novo* assembled into 67 million contigs for mosquito species identification and virus discovery.

**Fig. 1.**
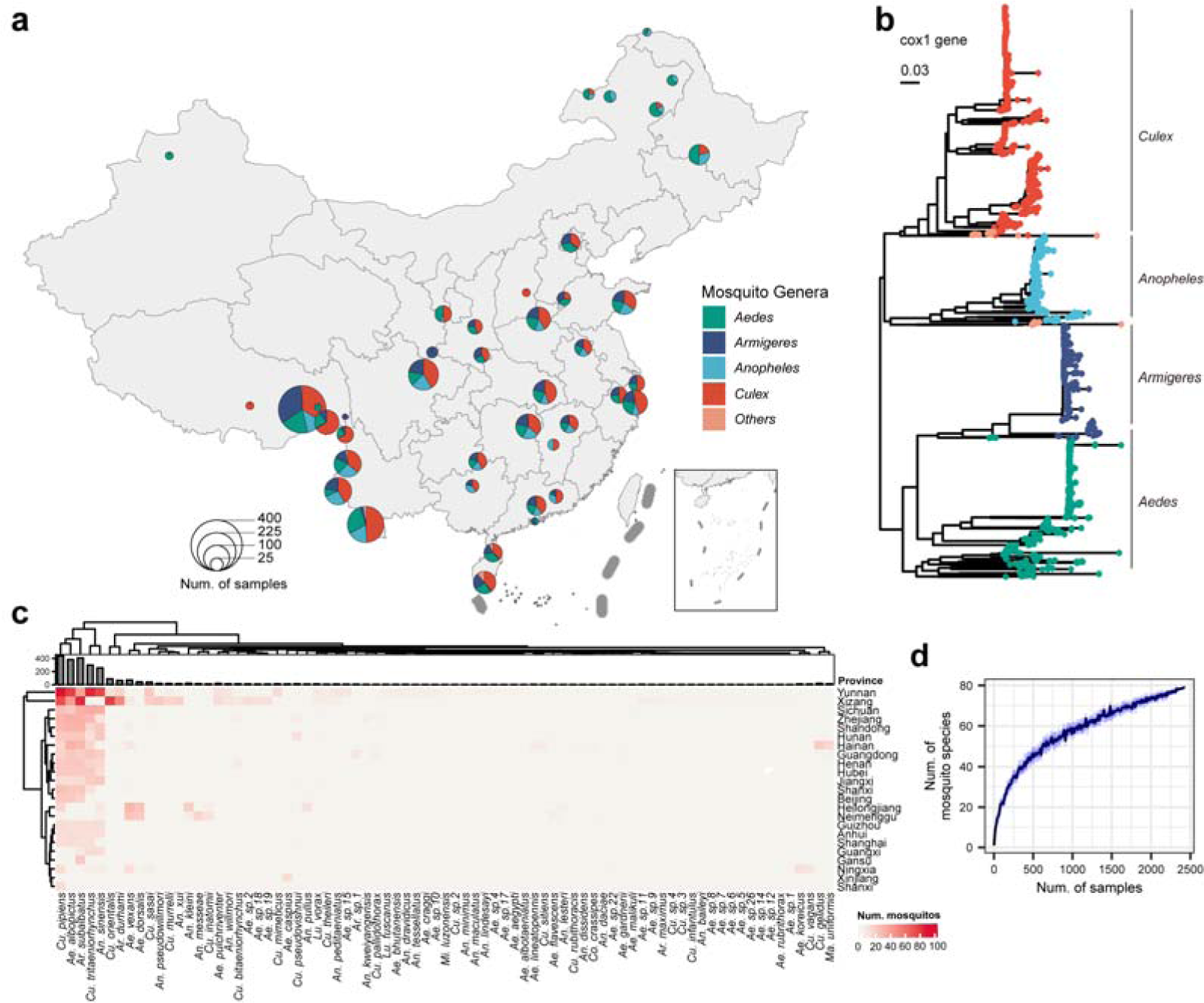
| Overview of the 2,438 mosquito individuals sampled across China. **(a)** Sample overview, showcasing the mosquito genera composition at each location. The pie chart area represents the number of sampled mosquitos. **(b)** Maximum likelihood phylogenetic tree depicting the relationships among the 2,438 mosquito individuals, constructed using the *cox1* gene. **(c) The composition of collected mosquito species in each province. (d)** Rarefaction curve displaying the richness of mosquito species, with the blue area indicating the 95% confidence interval.

Mosquito species identification was performed based on the mitochondrial gene *cox1*. This revealed 79 species belonging to eight genera (*Aedes*, *Armigeres*, *Anopheles*, *Culex*, *Mansonia*, *Mimomyia*, *Coquillettidia*, and *Lutzia*) (Fig. 1b). The most prevalent species includes well-known disease vectors: *Aedes albopictus* (n=383), *Armigeres subalbatus* (n=408), *Anopheles sinensis* (n=256), *Culex pipiens* (n=438), and *Culex tritaeniorhynchus* (n=298), which accounted for 73.1% of the samples (Fig. 1c; Fig. S1). These dominant species exhibited wide distribution across the country, spanning latitudes from 18° to 35° and longitudes from 26° to 35° (Fig. S2).

Viruses were identified using hallmark genes (e.g., the RNA-dependent RNA polymerase (RdRp) for RNA viruses and DNA polymerase for DNA viruses except the *Circoviridae* where Rep gene was used and the *Parvoviridae* where NS1 gene was used), compared against the NCBI non-redundant protein database. This process yielded a total of 205,032 viral contigs. After excluding retrotransposons, endogenous virus elements, and bacteriophage, we identified 564 distinct viral species (Supplemental Data 2). Of these, 393 are likely to infect mosquitos in contrast to other host taxa, and hereafter, define as being the core mosquito virome. The core virome, which comprises arthropod-borne arboviruses which may cause diseases in humans or other vertebrates (e.g., flaviviruses, alphaviruses, etc.), as well as insect-specific viruses that may directly affect mosquitos, is most likely to have a more important impact on human or animal (both vertebrates and invertebrates). Identification of mosquito-associated viruses was based on their close phylogenetic relationship to known mosquito-infecting viruses and their high viral abundance and prevalence in mosquitos (Fig. 2, Fig. S3-5; see also *Methods*). Through this process, viruses associated with other host taxa, such as fungi, protists or nematodes (parasites inside mosquitos or reagent contaminations), were excluded from downstream analyses.

**Fig. 2.**
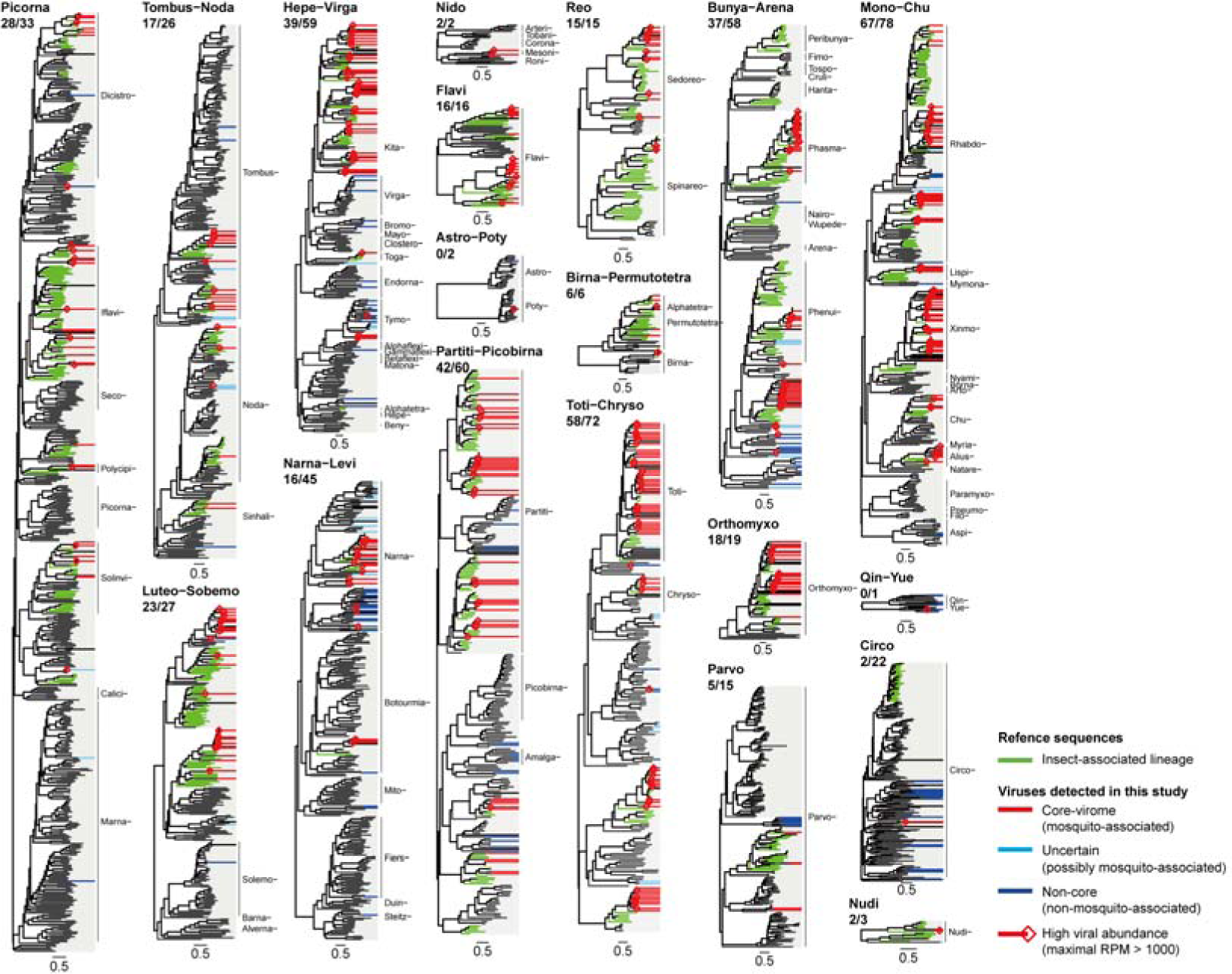
| Phylogenetic diversity of the 564 viruses discovered in this study and identification of the “core virome” of mosquitos. The phylogenetic trees were estimated using “hallmark” proteins of respective viral taxa. Trees of RNA viruses were estimated at the “super clade” level, while those of DNA virus were constructed at family level. The hallmark proteins utilized were the RNA-directed RNA replicase (RdRp) for all RNA viruses, Rep protein for the *Circoviridae*, NS1 protein for the *Parvoviridae*, and DNA polymerase for other DNA viruses. The number of mosquito-associated species (i.e., the core virome) and the total viral species of each viral super clade is shown below the superclade names (in the form of number of mosquito-associated species / total species). Viral families were shown indicated alongside the tree, and the suffix “-viridae” is omitted.

Our analysis revealed that individual mosquitos carried a median of two mosquito-associated virus species, with an interquartile range (IQR) of three and a maximum of 11. Viral RNA comprised 7.9×10^-6^%-84.5% of the total RNA (rRNA removed) within an individual mosquito, with an IQR of 0.01%-1.8%. A median of 85% of the total viral RNA within individuals belonged to mosquito-associated viruses (Fig. 3a). Rarefaction analysis indicated that the sequencing depth was adequate to reflect true diversity of viruses within individual mosquitos (Fig. S6).

**Fig. 3.**
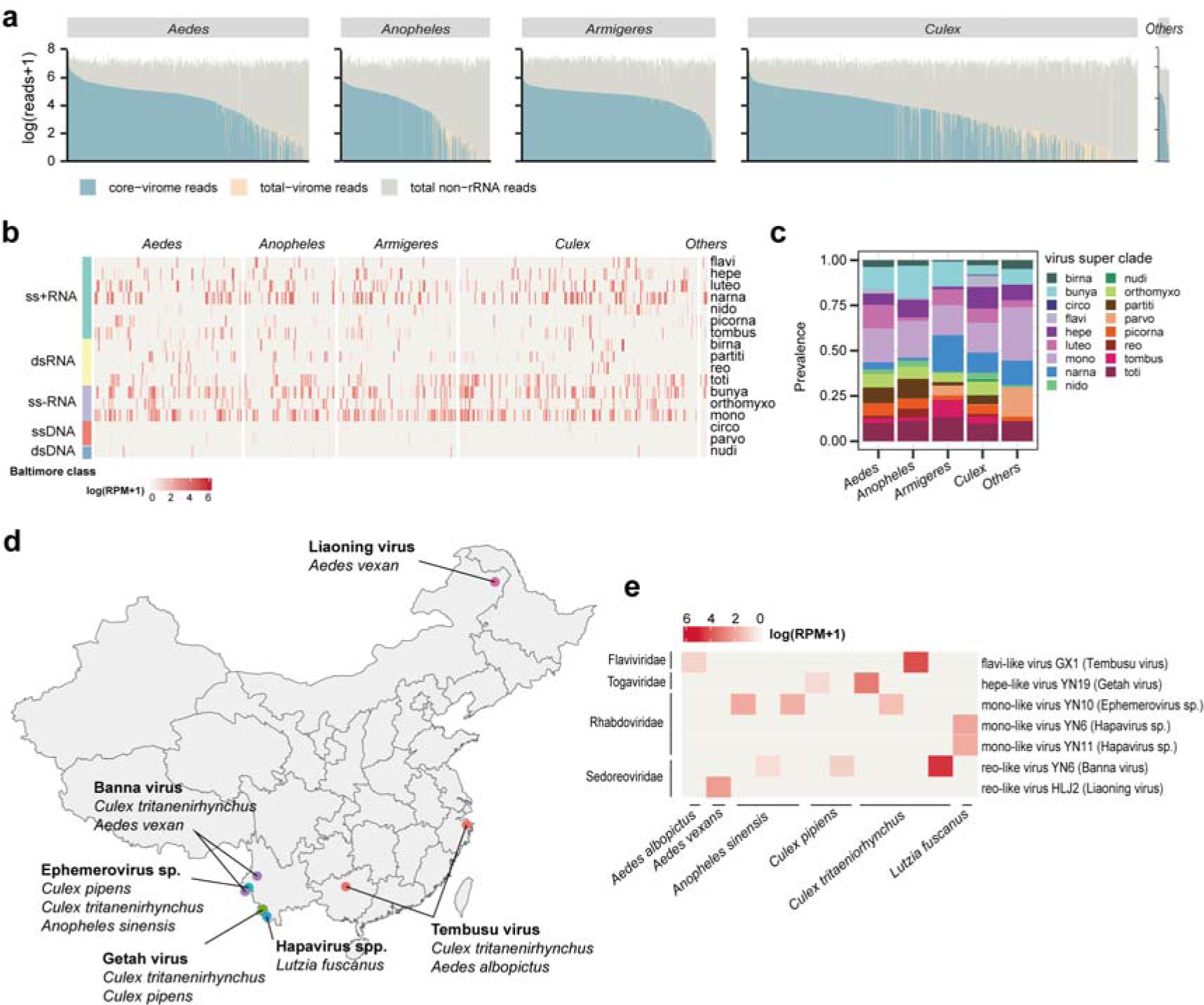
| Characterisation of individual mosquito viromes and the discovery of putative vertebrate-infecting arboviruses. **(a)** Sequencing depth and total viral abundance for each mosquito. **(b)** Composition of individual mosquito viromes. **(c)** A comparison of the prevalence of viral super clades within different mosquito genera. **(d)** Distribution of seven (putative) arboviruses. **(e)** Abundance of (putative) arboviruses within individual mosquitos.

A majority of the core virome had RNA genomes (384/393 species, 97.7%), which were classified into 16 previously established “super clades” that refer to clusters of related RNA viruses situated around the taxonomic levels of class or order^8^ (Fig. 2, Fig. 3b). Super clades were used because many of the viruses identified here are likely to represent novel or uncharacterised viral families, so that assignment of conventional taxonomic ranks may be imprecise. Among the 393 viral species, 249 (63.3%) were putative novel viral species, sharing less than 90% protein identity and 80% nucleotide identity with known viruses. Viruses with positive and negative single strand RNA genomes were the most prevalent (detected in 1370 (56.2%) and 1274 (52.3%) individuals respectively), followed by double-strand RNA (892, 36.6%), single-strand DNA (110, 4.5%), and double-strand DNA genomes (20, 0.8%) (Fig. 3bc). As for single virus species, most species were only detected in a few mosquito individuals (75% quantile: 12 individuals, 95% quantile: 59.35 individuals). In contrast, the top 2% prevalent virus species (such as Mosquito narna-like virus SC1 and Mosquito bunya-like virus SHX1) could be detected in more than 100 individuals, accounting for 40% ∼ 70% of individuals of a single mosquito species.

Overall, we identified four known and three potential arboviruses (Fig. 3de; Fig. S7). The four known arboviruses were Getah virus (genus *Alphavirus*), Tembusu virus (*Flavivirus*), Banna virus (*Seadornavirus*) and Liaoning virus (*Seadornavirus*), while the three putative arboviruses were Mosquito mono-like virus YN6 and YN11 (*Hapavirus*) and Mosquito mono-like virus YN10 (*Ephemerovirus*). The prevalence of these putative arboviruses was low, with detection only in 13 out of the 2,438 individuals. Most positive individuals were sampled in semi-natural habitats in southwestern China, where tropical forests were present (Fig. 3d). Despite of their low prevalence, these arboviruses can reach relatively high abundance within individuals (median RPM: 19 for arboviruses and 8.35 for all non-arboviruses), constituting up to 4.2% of the total non-ribosomal RNA and implying a high risk of transmission through a single bite (Fig. 3e).

### The Patterns and Drivers of Viral Diversity in Mosquitos

To investigate the environmental and host factors that shape viral species richness, we examined the effect of mosquito species identity, land-use characteristics and various climatic variables using generalized linear models (Fig. 4). The best model structures were selected by examining all combinations of variables based on the AIC criterion. This revealed that for the best model, mosquito species identity explained the most deviance (17%), followed by climate (4%) then by land-use (1%). Most deviance remained unexplained (78%) (Fig. 4c). As for individual variables, mosquito species identity, mammal richness, and climate were consistently support by the seven supported models (ΔAIC<2) (Fig. 4b).

**Fig. 4.**
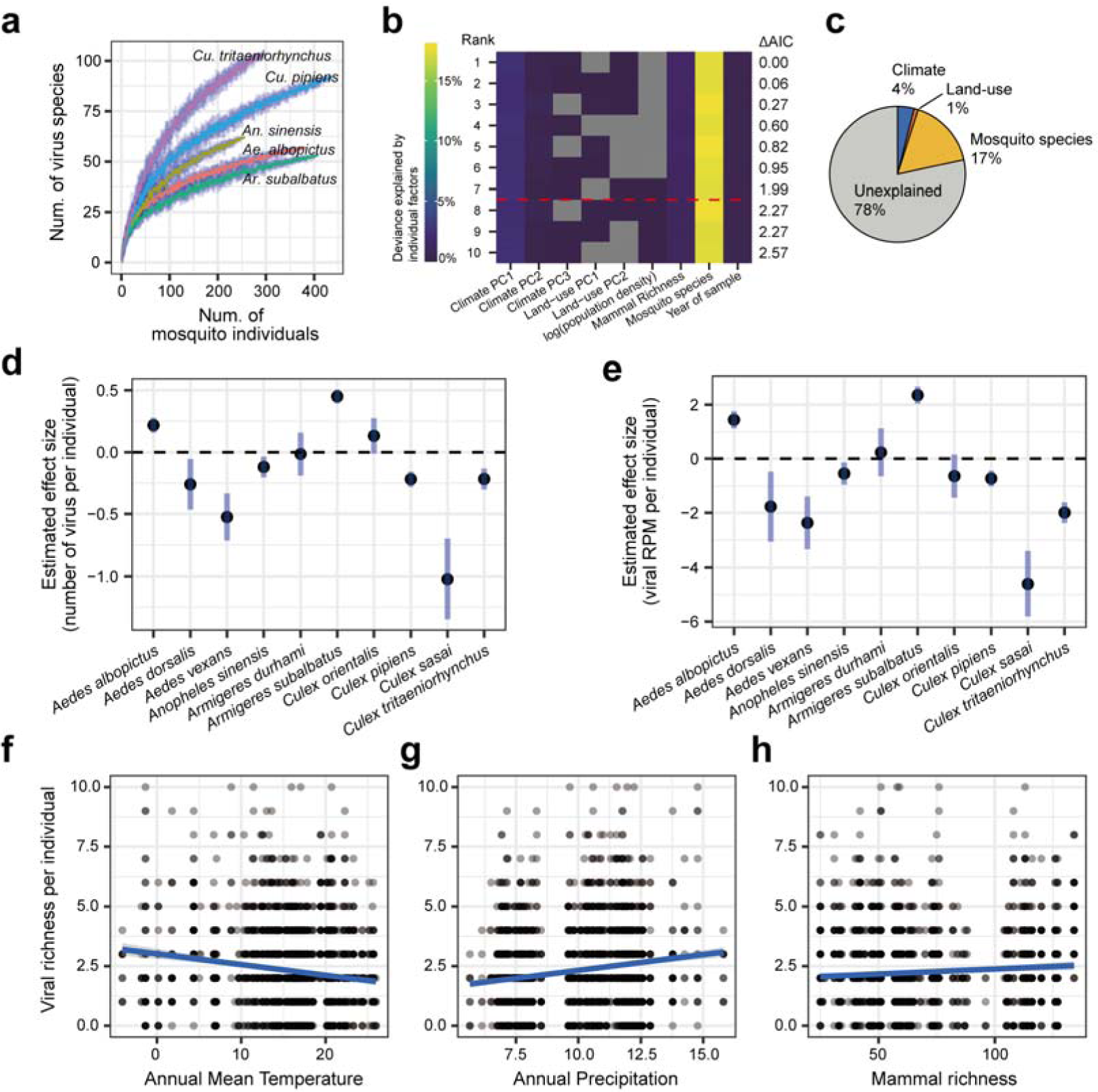
| Environmental and host drivers of viral diversity. **(a)** Rarefaction curves for viral richness discovered in the five dominant mosquito species. The blue areas represent the 95% CI. **(b)** Relative effects of mosquito species, climate, and land-use characteristics on viral richness per individual mosquito. The relative effect of factors was quantified by explained deviance in generalized linear models. The top ten models selected by Akaike information criterion (AIC) are shown. The red dashed line indicates the four most supported models (ΔAIC < 2). “PCs” abbreviate principal components. **(c)** Relative contribution of mosquito species, climate, and land-use characteristics to viral richness. **(d)** Model estimations of the effect of mosquito species on viral richness per individual. Error bars indicate 95% confidence intervals (CI). **(e)** Model estimations of the effect of mosquito species on total viral abundance per individual. Error bars indicate 95% CI. **(f)** Relationship between mean annual temperature and viral richness per individual. **(g)** Relationship between annual precipitation and viral richness per individual. **(h)** Relationship between mammal richness and viral richness per individual.

Notably, the distribution of viral richness was uneven among mosquito species (Fig. 4d). As for the five dominant mosquito species, *Ar. subalbatus* carried the highest number of viral species per individual (3.14±1.63 species, mean±S.D.), followed by *Ae. albopictus* (2.60±1.76), *An. sinensis* (1.94±1.78), *Cu. pipiens* (1.84±1.65) and *Cu. tritaeniorhynchus* (1.68±1.61). The total viral abundance within individual mosquitos also varied among mosquito species (Fig. 4e). Viral species richness per individual mosquito was positively associated with mammal species richness and mean annual precipitation, while negatively associated with mean annual temperature (Fig. 4f-h). Similar trends were observed for total viral species richness within mosquito populations (Fig. S8; a population represents all individuals of a mosquito species within 100 km diameter centred around a sample location).

### The Patterns and Drivers of Virome Composition in Mosquitos

We next investigated the environmental and host factors that shape virome composition at both individual and population scales using distance-based redundancy analysis and forward model selection (Table S1). At the population scale, model selection indicated mosquito species as the only significant factor shaping virome composition, explaining 33.9% of the total variance. At the individual scale, mosquito species remained the most important, explaining 11% of the total variance, but was not the only factor shaping virome composition. Climate (three climate principal components) and land-use (two anthrome principal components and mammal richness) factors explained 2%, and the rest part of variance (87%) remained unexplained.

These results indicate that the mosquito viromes were distinct among different mosquito taxa at both individual and population scales, which was then visualised using t-SNE (t-distributed stochastic neighbour embedding) (Fig. 5a). Interestingly, 192 viral species were still shared among multiple mosquito species, including 123 species shared among genera (Fig. 5b-d).

**Fig. 5.**
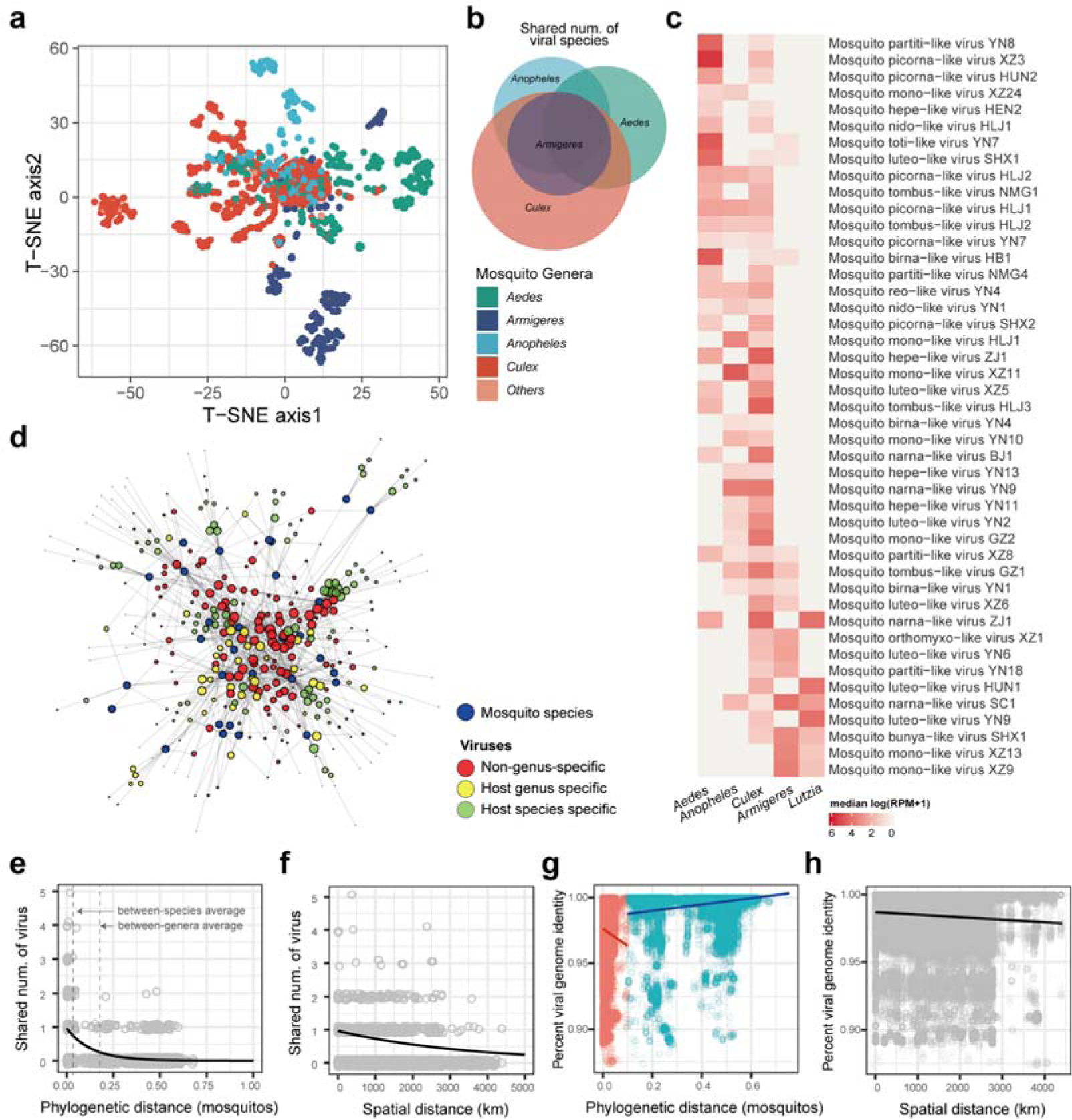
| The composition of individual mosquito viromes and the drivers of viral sharing among mosquitos. **(a)** The t-SNE ordination of individual mosquito viromes. Colours represent mosquito genera. **(b)** Venn diagram showing the number of shared and unique viral species identified in each mosquito genus. **(c)** The abundance of viruses that were shared among different genera. **(d)** The virus sharing network among mosquito species. For visual clarity, only the top 30 abundant mosquito species and their viromes were shown. **(e)** The effect of mosquito phylogeny on viral sharing between pairs of mosquito individuals. The two dashed lines indicate average phylogenetic distance between species and between genera respectively. **(f)** The effect of spatial distance on viral sharing between pairs of mosquito individuals. **(g)** The relationship between mosquito phylogeny and viral genome similarity. The two dashed lines indicate average phylogenetic distance between species and between genera respectively. Smoothed lines are interpolated using cubic splines. **(h)** Distance decay of viral genome similarity.

To disentangle the potential drivers of the prevalent virus sharing across species and genera, we analysed the number of shared viral species between each pair of mosquito individuals. Using a generalised linear model, we found that the number of shared viral species decreased sharply with increasing phylogenetic distance (branch length of the *cox1* gene phylogeny) between individual mosquitos (Fig. 5e), after accounting for the effects of covariates (sampling date and spatial distance). Interestingly, the decrease was less pronounced with increasing spatial distance (Fig. 5f). Even when two individual mosquitos were separated by distance of thousands of kilometres, they could still share the same viral species (Fig. 5f).

We also analysed patterns of virus-sharing among mosquito individuals at intra-specific scale of each virus species. We compared the genome identity of variants from each viral species (Fig. 5gh). This revealed that the similarity between viral genomes declined as the phylogenetic distance of mosquito individuals increased if the phylogenetic distance was smaller than 0.167 (i.e., the average phylogenetic distance between mosquito genera), indicative of co-divergence between viruses and their hosts. However, if the phylogenetic distance exceeded 0.167, the similarity between viral genomes remained high (95% quantile: 95.7%) and displayed no significant trends with phylogenetic distance of mosquito individuals. This suggests that mechanisms of host adaptation may differ when a virus is transmitted within the same host species compared to the case when it is transmitted among different species or genera.

### Host Specificity of Mosquito-Associated Viruses

Based on these data, we hypothesized that mosquitos harbour both generalist and specialist viruses. Although some viruses can be detected in multiple host species, the viral abundance can be low in these cases, suggesting transient spillover (Fig. 6b). To examine host specificity more robustly we quantified the degree of virus-host co-divergence by correlating phylogenetic distance (i.e., the sum of branch lengths on the maximum likelihood phylogenetic trees) among variants of each viral species with that of their hosts, using partial Mantel tests (the effects of spatial distance among hosts were considered simultaneously). Only viruses detected in more than five individuals were included. This analysis revealed a spectrum of host generalist to specialist viruses (Fig. 6). In particular, 23 of 237 mosquito-associated virus species exhibited significant co-divergence with their hosts after removal of the effect of spatial distance (partial Mantel tests, r > 0 and p < 0.05).

**Fig. 6.**
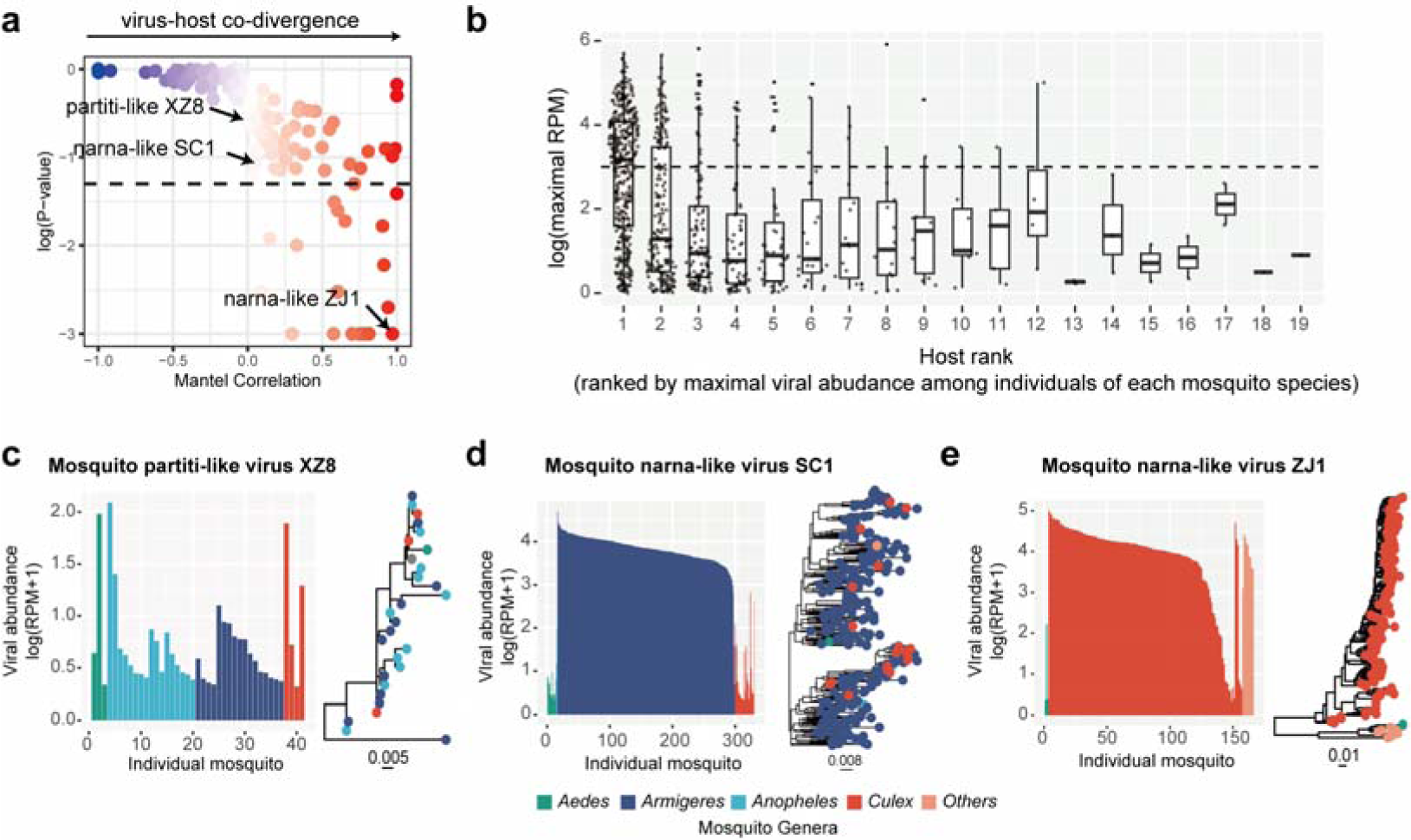
| Host specificity of mosquito-associated viruses. **(a)**. The degree of virus-host co-divergence was measured using the Mantel correlation (Spearman) between the phylogenetic distance matrix of viruses and their corresponding mosquito hosts. Colours indicate correlation coefficient (the same as the X axis). **(b)** Comparison of the viral abundance of each viral species in its major and minor hosts (ranked by maximal viral abundance among all individuals of a mosquito species). The dashed line marks the position where RPM=1000, indicating high viral abundance. **(c, d and e)** Abundance of three example viral species Mosquito partiti-like virus XZ8, Mosquito narna-like virus SC1 and Mosquito narna-like virus ZJ1 within individual mosquitos, along with their respective phylogenetic trees. Phylogenetic trees were estimated using whole genome sequences and the maximal-likelihood method.

The differences in host specificity are illustrated by three examples. The first – Mosquito partiti-like virus XZ8 – appeared to be a generalist virus as it was found in four different mosquito genera and did not exhibit co-divergence with its host species (Fig. 6c). In addition, its abundance within each host was relatively low. In contrast, Mosquito narna-like virus ZJ1 appeared to be a specialist virus. Although it was detected in three genera, it showed significant co-divergence with its hosts. Each host genus formed a monophyletic group, and the viral abundance was high (Fig. 6e). Similarly, Mosquito narna-like virus SC1 seemed to be a specialist virus associated with the mosquito genus *Armigeres*, although strains infecting other host genera were observed in the Armigeres-infecting clade, suggesting frequent host jumping events (Fig. 6d). Thus, the host specificity of viruses is likely to be a continuum from generalist to specialist.

### Linking Virus Biogeography with Mosquito Phylogeography

Our results revealed that even when two individual mosquitos were separated by thousands of kilometres, they could still share the same viral species, indicative of wide distribution of some viral species (Fig. 5f). For example, 76 of the 393 mosquito-associated viruses were widespread across China (median distance between positive individuals >1000km, indicating large spatial converge), with the remainder being more localised (median distance <1000km) (Supplementary Data 2). Viruses detected in multiple host species have a wider distribution range compared to those associated with a single mosquito species, after the effect of sample size bias (i.e., number of positive individuals of each virus may vary) was considered using multiple linear regression (Fig. S9).

We next explored the intra-specific genetic structure of each virus species throughout space. We quantified how viruses diverged as spatial distance increases, by correlating phylogenetic distance among variants of each virus species with the spatial distance between them, using partial Mantel tests (to remove the effect of host phylogeny). This analysis revealed 79 virus species that displayed significant genetic structure in space (partial Mantel test, r > 0 and p < 0.05; Fig. S10, Supplemental Data 2). Viruses that were genetically divergent in space did not necessarily co-diverge with their hosts (χ^2^ test, p =0.003; Fig. S10), such that the spatial structure of viruses was unlikely to be driven by co-divergence with their hosts. Of note, 65 of the 76 viruses considered to be widespread did not show significant spatial genetic structure. In addition, genetic divergence of those viruses remained low across long distances (whole genome substitution rate was 0.025 on median, IQR=0.027, inferred from maximal likelihood tree), tentatively suggesting rapid spread in their recent history.

To determine how the dispersal of mosquitos contributed to the biogeographic patterns observed in mosquito viromes we correlated the number of shared virus species between mosquito populations with the phylogenetic distance of mosquitos among populations and spatial distance (Fig. 7a). The results indicated that the number of shared viruses between populations declined sharply with minimal *cox1* distance among host populations. In contrast, there was a much weaker correlation with spatial distance (Fig. S11). This can be visualised by comparing the networks of shared number of viruses among populations to the networks of host similarity (Fig. 7a). Although some mosquito populations are spatially close, the number of shared viruses between them is primarily determined by the genetic distance of mosquitos, suggesting the potential importance of long-range host migrations to virus dispersal.

**Fig. 7.**
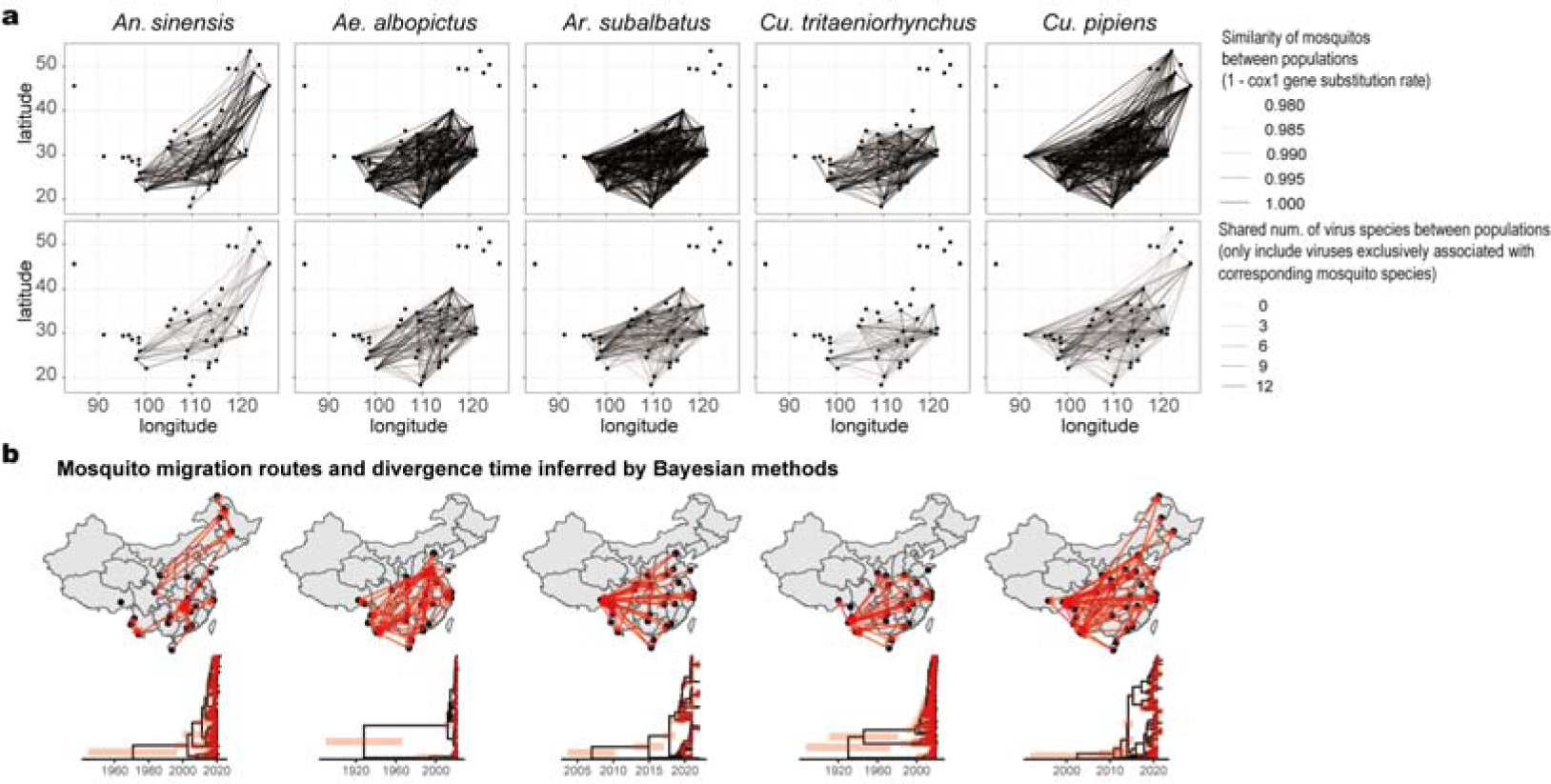
| Linking virus biogeography with mosquito phylogeography. **(a)** The networks of similarity of mosquitos and virome connectivity among mosquito populations. The first row visualises the minimal phylogenetic distance among mosquitos between pairs of mosquito populations. The darker lines indicate lower phylogenetic distance. The second row visualises the connectivity of virome between pairs of mosquito populations. The darker lines indicates that more virus species are shared. **(b)** The migration routes and divergence time estimated by Bayesian methods. The first row displays the reconstructed routes of historical migration of mosquitos. All shown routes were supported by the Bayesian stochastic search variable selection (BSSVS), with Bayes Factors (BF) greater than three. The second row shows the estimated divergence time of mosquitos. Molecular clocks were calibrated using four internal calibration points based on fossil records of oldest Diptera species (calibration point: Diptera), the earliest fossil mosquito (Culicidae) and the earliest Culicinae subfamily record. Red bars represent 95% HPD.

To analyse this further, we inferred the phylogeographic history of the five most dominant mosquito species using the *cox1* gene. Migration routes were then inferenced using a Bayesian stochastic search variable selection (BSSVS) process, and divergence times were estimated using a molecular clock calibrated using fossil evidence^22–24^ (Table S3). These data showed that the divergence and migration of mosquitos across the country likely only occurred recently (Fig. 7b). Together, rhese results suggest that the long-distance dispersal of mosquitos may have contributed to the formation of large-scale biogeography of mosquito viromes.

## DISCUSSION

We collected 2,438 individual mosquitos from diverse habitats across China and characterized the virome of individual mosquitos by meta-transcriptomic sequencing. To the best of our knowledge, there have been only five studies that have characterised the virome of individual insects^11–15^, all restricted to small areas and involved limited individuals. Other metagenomic studies, in which samples were pooled by species or locations, were unable to reveal the patterns and drivers of viral diversity at individual scale^7–10^. Our expansive data set of individual viromes of over 2,000 mosquitos linked the viral diversity within individual insects with total diversity at country scale, providing unique insights into the general ecology of the viromes of insect vectors (e.g., virus prevalence, distribution, drivers of virome composition, host specificity).

Our results revealed a very low prevalence of vertebrate-associated arboviruses within mosquitos. By sequencing over 2,000 mosquito individuals in the most populated areas of China, we only detected 13 individual mosquitos that were positive for any known or putative arboviruses. This is despite the fact that we studied well-known disease vectors such as *Ae. albopictus* (Asian tiger mosquito) and *Cu. pipiens* (common house mosquito), and the corresponding sample size for these species are large (383 and 438 individuals respectively). This result generally aligns with findings of previous metagenomic studies of vector arthropods, such as mosquitos^11^ and ticks^7^.

Notably, the arboviruses detected were concentrated in semi-natural habitats nearby forests, suggesting that their prevalence is uneven among locations or habitats. While it is well-documented that the prevalence of mosquitos can differ between regions or habitats in the case of some arboviruses (such as West Nile virus^25,26^ and Japanese encephalitis virus^27^), such evidence is lacking for relatively understudied arboviruses, as well as for newly discovered putative arboviruses. Our metagenomic data therefore provides powerful new evidence to support an uneven prevalence of arboviruses. Such distribution patterns also tentatively suggests that wild animals might be involved in the transmission of these viruses. Many other well-studied arboviruses are known to be maintained in populations of wild animals through enzootic transmission cycles^28^, including West Nile virus and Japanese encephalitis virus. As such, these results highlight the importance of intensifying pathogen surveillance efforts in these semi-natural habitats, as they may act as the frontline for the spillover of novel or neglected/understudied arboviruses to humans.

We also identified hotspot species and locations of overall viral diversity, which largely coincided with arbovirus hotspots. We detected significant variation in the diversity of individual viromes across mosquito species. This implies that there are hotspot species of viral diversity. Although previous studies have also reported that the diversity and composition of virome differ among mosquito species^10,12^, they were unable to identify hotspot species due to the insufficient sample size per species. Our results offer novel evidence for the existence of diversity hotspot species. While comparative immunological studies have associated various anti-virus immune genes with the vector competence of a few well-studied arboviruses^29–31^, it remains largely elusive whether such immunologic variation will affect the total viral diversity. We hypothesize that such variation among mosquito species reflects their carrying capacity for viruses and is ultimately indicative of the total investment in mosquito immunity. These differences could result in “hotspot” species that harbour more viruses and exhibit higher viral loads, thereby potentially posing a higher risk of arbovirus emergence.

Our findings also suggested that areas with high mammal richness, relatively low temperatures, and high precipitation may act as hotspot areas for viral diversity. These diversity hotspot areas correspond to previously described distributions of arboviruses, further emphasizing the need for surveillance efforts in these areas. Notably, a considerable portion of the deviance in virus diversity remains unexplained. This could be due to unmeasured environmental or host covariates or pure stochasticity^32^. For example, intraspecific trait variations (e.g., immune phenotypes) among mosquitos could be an important but unmeasured factor. Clearly, the relative contribution of deterministic and stochastic processes in shaping the diversity and composition of individual mosquito viromes remains unknown, and further studies are warranted.

Remarkably, our data uncovered a substantial number of viruses that were shared among different mosquito species or genera, broadening our understanding of the host specificity of insect-associated viruses. While our findings indicate that phylogenetic distance among mosquitos constrained virus transmission, there are cases of generalist viruses that can infect multiple mosquito species. Hence, counter to earlier ideas, virome compositions are not completely distinct among insect species^7,10,12^. Notably, our results also imply that the host specificity of mosquito-associated viruses exists on a spectrum rather than adhering to a rigid “generalist-specialist” dichotomy^28,33^. Some viruses appear to be predominantly associated with particular mosquito species, although sporadic host switching events are frequent, such as the case of Mosquito narna-like virus SC1. This phenomenon of frequent host switching has also been observed in rabies viruses of bats^34^, and in recent metagenomics studies in mammals^6^ and ticks^7^. The frequent spillover events, along with the frequent co-infection of viruses within individual mosquitos, are likely to contribute to increased genetic diversity among viruses circulating in mosquito communities. Furthermore, studying the differences in the determinants of host competence between generalist and specialist viruses holds considerable value. For example, some studies have suggested that many arboviruses are generalist in nature^7,28^, and understanding this may be important for disease control. Therefore, understanding the general principles underlying virus-host specificity and the determinants of host competence is a promising avenue for arbovirus prevention.

We also detected a wide array of virus species that were widespread across China as a whole, with highly similar genomes throughout. This phenomenon may be attributed to recent long-distance dispersal events. Although it is presumed that the rapid spread of pathogens through mosquito flight should be limited due to the weak flight ability of mosquitos, our analyses of cox1 gene similarity and phylogeographic reconstructions suggest that the five dominant mosquito species have likely spread throughout the country in recent history, which is consistent with some previous evidence^35^. As a consequence, the dispersal of mosquitos may be an important factor that shapes the biogeography of mosquito-associated viruses. Previous studies have highlighted infected travellers, rather than mosquitos themselves, as the main driving force behind virus spread^36,37^. Our results emphasizes that the movement of mosquitos, potentially facilitated by human activities, should not be overlooked. Human activities, such as transportation (e.g., the trade of used tires, which was a major route for the spread of Asian tiger mosquitos^38^), may also facilitate mosquito dispersal. Previous studies have also indicated the potential for wind-facilitated seasonal migration of mosquitos^39^ and other insects^40^. For instance, *Anopheles* mosquitos in Africa are known to migrate seasonally at high altitudes (>150m) by wind, rapidly transmitting malaria to distant areas^39^. When combined with our data, these suggest the importance of mosquito spread in virus dissemination.

In summary, our unique data set of individual mosquito viromes provides important insights into the micro- and macro-diversity of viromes associated with vector insects. We identified potential hotspot locations and species of viral diversity and potential vector-borne diseases. These findings highlight the need for enhanced virus surveillance in these potential high-risk areas and among these species. Additionally, we have demonstrated that the composition of individual viromes is strongly influenced by host phylogeny, although generalist viruses that are shared among different host species/genera are prevalent. Finally, we have shown that the often-overlooked long-range dispersal of insect host may play a crucial role in shaping virus distribution, alerting us to consider pathways of vector insect dispersal such as transportation and wind-facilitated spread.

## METHODS

### Sample collection

A total of 2438 adult mosquitos were collected between 2018 to 2021 in China. The sample locations involved 23 Chinese provinces, including Tibet, Xinjiang, Yunnan and Inner Mongolia Provinces, spanning approximately 4000 km both in latitude and longitude (Fig. 1; Supplemental Data S1). Mosquitos were collected using CO_2_ traps (dry ice) that were set at each location for approximately 12 h overnight. Each trap was baited with dry ice to attract mosquitos. Upon trap collection, the mosquitos were collected and preserve in dry ice. Mosquitos were then placed in labelled vials and left on dry ice until they were returned to the laboratory, where the samples were placed in a −80°C freezer until RNA extraction. Mosquito species identification was initially carried out by experienced field biologists using taxonomic keys and dissecting microscopes on cold tables and was later verified by analysis of the cytochrome c oxidase subunit I (cox1) gene (Fig. 1; Supplemental Data S1).

### Sample Processing and Sequencing

RNA extraction and sequencing were carried out on each mosquito individual (Supplemental Data S1). Prior to homogenization, each mosquito pool was washed three times with 1 ml of a sterile RNA- and DNA-free phosphate-buffered saline (PBS) solution (Gibco) to remove external microbes. The samples were then homogenized in 600μl of lysis buffer by using a TissueRuptor instrument (Qiagen). Total RNA was extracted by using a RNeasy Plus minikit according to the manufacturer’s instructions. The quality of the extracted RNA was evaluated by using an Agilent 2100 bioanalyzer (Agilent Technologies). All extractions performed in this study had an RNA integrity number (RIN) of >8.7.

The sequencing libraries were prepared using the MGIEasy RNA Library Prep Kit V3.0. Briefly, the RNA was fragmented, reverse transcribed and synthesised into double-stranded cDNA. The Unique Dual Indexed cDNA was circulated, and rolling-circle replicated into DNA nanoball (DNB)-based libraries. The constructed libraries were sequenced on the DNBSEQ T series platform (MGI, Shenzhen, China) to generate meta-transcriptomic data of 150-bp paired-end reads.

### Virus Discovery

For each sample, reads from ribosomal RNA (rRNA) were removed using URMAP^41^ (version 1.0.1480), and then by mapping quality-controlled raw reads against the SILVA^42^ rRNA database with Bowtie2^43^. Adapters and low-quality reads were removed using fastp^44^ (version 0.20.1). The reads with duplicates and low complexity were removed using SOAPnuke^45^ (version 2.1.5) and PRINSEQ++^46^ (version 1.2), respectively. The resulting clean non-rRNA reads were assembled into contigs using MEGAHIT^47^ (version 1.2.8). The assembled contigs were searched against the NCBI nr database using DIAMOND (version 2.0.15). The e-value was set at 1e-5 to achieve high sensitivity while reducing false positives. We roughly classify the contigs by kingdom based on the search, and we extracted all viral contigs. Viruses were further confirmed by checking the existence of hallmark genes (i.e., RdRp for RNA viruses, NS1 for *Parvoviridae*, and DNApol for other DNA viruses). We removed viral contigs that were less than 600bp in length, and we also checked the domain completeness for each RNA virus (at least one conserved motif of RdRp should exist, checked manually by performing multi-sequence alignments). Viral contigs with unassembled overlaps were merged using SeqMan in the Lasergene software package (version 7.1). To confirm the assembly results, reads were mapped back to the virus genomes with Bowtie2.

### RNA Quantification

To quantify the abundance of RNA genomes (RNA viruses) or transcripts (DNA viruses), we estimated the percentage of total reads that mapped to target genomes or genes. The sequences used for mapping involved viral genomes and mosquito marker gene (cox1) mentioned above. Mapping was performed using Bowtie2. We used number of reads mapped per million non-rRNA reads (RPM) to represent viral abundance.

### Virus Discovery Quality Control

To ensure that the viruses we detected in mosquito individuals were not artifacts, we applied multiple quality control measures. First, we confirmed that the individual mosquitos sequenced were not contaminated by other individuals or species by examining the polymorphism of the mitochondrial cox1 gene. We mapped sequencing reads to the cox1 gene and determined whether there were polymorphic sites in the mapped regions. As there was only one individual sequenced in each library, there should be no polymorphic sites in the cox1 gene (with the exception of sequencing error). We confirmed that the frequency of single nucleotide substitution in the cox1 gene was less than 1% for each of the 2438 individuals, supporting that our samples and sequencing process were not contaminated.

We then performed two rounds of filtrations based on viral abundance. First, we removed false positives due to index-hopping that occurs during high-throughput sequencing when reads from one sample are erroneously assigned to another sample. We minimized the effect of index-hopping by applying a read abundance threshold relative to the maximal read abundance within each sequencing lane. If the total read count of a specific virus in a specific library of is less than 0.1% of the highest read count for that virus within the same sequencing lane, then it is considered as a false-positive due to index-hopping. In addition, we excluded those viruses at very low abundance (RPM < 1) and at low genome coverage (coverage < 300 base-pairs, i.e., the length of two reads), which were also likely to be false-positives. These thresholds have been validated in our previous publications deploying RT-PCR re-confirmation^6,48^.

### Phylogenetic Analyses

To determine the evolutionary history of the newly discovered viruses, we estimated viral phylogenies using the amino acid sequences of the viral hallmark proteins (i.e., RNA-dependent RNA polymerase for RNA viruses, DNA polymerase for DNA viruses, except Rep protein for the *Circoviridae* and NS1 protein for the *Parvoviridae*). For comparison, we included previously reported viral hallmark protein of each relevant phylogenetic groups (RNA viruses were grouped by super clades^8^, and DNA viruses were grouped by viral families). This included all the previously described mosquito viruses within these groups. Within each group, the hallmark proteins were aligned by using the E-INS-i algorithm implemented in MAFFT^49^ (version 7). Ambiguously aligned regions were removed using TrimAl^50^. The best-fit model of amino acid substitutions was determined busing jModelTest^51^. Phylogenetic trees were inferred by using the maximum likelihood (ML) method implemented in PhyML^52^ version 3.0, utilizing the best-fit substitution model and the Subtree Pruning and Regrafting (SPR) branch-swapping algorithm. Support for individual nodes on the phylogenetic tree was assessed by using an approximate likelihood ratio test (aLRT) with the Shimodaira-Hasegawa-like procedure as implemented in PhyML.

### Identification of the Core Mosquito Virome

The core virome in this study refers to viruses associated with mosquitos, as distinguished from viruses associated with other host taxa such as protists, nematodes, fungi, bacteria amongst others. The identification of the core virome was based on four key aspects: (i) similarity to viruses known to infect specific hosts, (ii) viral abundance within individual mosquitos, (iii) phylogenetic position on the tree, and (iv) co-occurrence with other microbes within the same individual. To accomplish this, we gathered genome/protein sequences and corresponding host information for all viruses listed in the ICTV master species list and the Virus-Host Database (Fig. S3-4). Using a BLASTP search, we compared the hallmark proteins (RNA viruses: RdRp, *Circoviridae*: Rep, *Parvoviridae*: NS1, other DNA viruses: DNApol) of the viruses detected in our study against these databases. We assigned host annotations to each virus based on the best BLASTP hit, considering hits with an e-value greater than 10^-5^ and a percentage identity above 40%. The thresholds were benchmarked using the above two databases to have achieved low false-positive rate and descent recall (Fig. S5).

In addition, viruses with a maximum abundance exceeding 1000 RPM among all positive individuals were roughly categorized as “mosquito-associated” (i.e., part of the core virome). We also manually inspected the phylogenetic tree (see above, *Phylogenetic Analysis*) to determine whether the viruses belonged to known mosquito or insect-associated lineages. If a virus did not belong to any known insect-associated lineages but co-occurred with microbes such as protists or fungi in the corresponding individuals, we classified it as non-mosquito-associated (i.e., non-core virome). In cases where a virus showed distant evolutionary relationship to any known viruses, exhibited paraphyly with known lineages, and had low abundance, we assigned it as “uncertain”.

### Inference of Mosquito Phylogeography

The phylogeographic history of the five dominant mosquito species was estimated using the BEAST^53^ software (version 1.10.4). Accordingly, we aligned the nucleotide sequences of the mitochondrial cox1 gene for each mosquito species using MAFFT. The resulting alignment files were then provided as inputs to BEAST. For the substitution model selection, we employed jModelTest and determined that the GTR+I+F+G model was the optimal choice. Two independent runs were conducted for each mosquito species, consisting of 50 million steps, with the results from the two runs were combined later. Sampling occurred every 1000 steps, with the initial 20% of samples were discarded as burn-in. Coalescent tree priors were set to a constant size model. To calibrate the molecular clock, we utilized three internal calibration points based on fossil records (Table S3) and incorporated tip dates as well. To account for rate variation among branches, we employed a relaxed molecular clock with an uncorrelated lognormal distribution. Convergence of the chains was assessed using TRACER^54^ v1.7 to ensure that the effective sample sizes (ESS) exceeded 200, indicating sufficient sampling. The Bayesian stochastic search variable selection (BSSVS) procedure was applied to estimate the historical migration of mosquitos among different sampling locations. Transition rates among locations and Bayesian factors were estimated, with transition rates supported by a Bayes factor (BF) greater than 3 considered to have significant support.

### Collection of Environmental Data

To examine how the diversity and composition of mosquito viromes are shaped by environmental factors, we collected climate and land-use data for each sample location from open data sources. Climate data were collected from the WorldClim2 database^55^. We extracted all the 19 “bioclimatic variables” commonly used in species distribution modeling and related ecological modeling techniques^56^. The definitions of these 19 variables can be found at https://www.worldclim.org/data/bioclim.html. To deal with co-linearity between climate variables, we conducted principal component analyses (PCA), and the first three principal components (i.e., CPC1, 2, and 3) were used to represent climate in all downstream statistical analyses. These principal components explained 52.4%, 27.1%, and 10.2% of the total variance respectively (sum up to 89.7%).

Land-use data were retrieved from the HYDE 3.2 database^57^ (spatial resolution: 5 arc minutes). We used the anthrome (i.e., anthropogenic biomes) classification system (21 classes in total) for land-use characterization. For robustness, we calculated the percentage of area classified as each class within 100 km diameter centered around each sample location, instead of using the anthrome classification exactly at each location. To remove co-linearity, we also calculated principal components for all anthrome variables. The first two components were used (referred to as APC1 and 2), and they explained 56.8% and 43.2% of variance respectively (sum up to approximately 100%). We also included the human population density in the year of 2017 from HYDE 3.2 database, and we retrieved mammal richness data (spatial resolution: 30*30 km) from the IUCN mammal richness database (version 2022-2).

### Statistical Methods

All statistical analyses were conducted using R version 4.2.0.

#### Estimating Alpha-Diversity in Individual Mosquito Viromes

To examine the influence of environmental and host factors on viral species richness, we employed generalized linear models with a Poisson distribution and log link function. The factors investigated included mosquito species identity, land-use characteristics, and climatic variables. Land-use characteristics comprised two principal components (APC1 and 2), log-transformed population density, and mammal richness. Climate conditions were represented by three principal components (CPC1/2/3). Model selection was based on the Akaike information criterion (AIC), assessing all combinations of variables using the MuMln package in R. The proportion of deviance explained by each variable was estimated by comparing the deviance explained by the full model to that explained by a model with the specific variable removed.

#### Composition and Connectivity of Individual Mosquito Viromes

The composition of individual mosquito viromes was visualized using the t-SNE method. We assessed the differences in virome compositions among mosquito genera through permutational multivariate analysis of variance (PERMANOVA) based on Jaccard distance. Additionally, the virus-sharing network was visualized using the igraph package in R to illustrate virome connectivity.

We also analyzed the effect of host phylogenetic distance and spatial distance on the pattern of virus-sharing among individuals. We employed generalized linear regression (Poisson regression) to examine the influence of host phylogeny and spatial distance on the number of shared virus species between pairs of mosquito individuals. To account for potential confounding effects, sampling date was included as a covariate in the analysis. To support the results of Poisson regression, we also conducted partial Mantel tests to correlate the number of shared-virus with phylogenetic distance among hosts, and spatial distance, while considering sampling date as a covariate. As the variables were not normally distributed, Spearman correlation coefficient were employed the partial Mantel tests.

#### Analysis of Host Specificity of Mosquito-Associated Viruses

We investigated the extent of co-divergence between viruses with their mosquito hosts, as well as the degree of virus divergence with increasing spatial distance. For each virus species detected in more than five positive individuals, we obtained consensus sequences within each positive mosquito individual by mapping RNA reads to its representative genome using bowtie2. The genome sequences were aligned using MAFFT, using the L-INS-I algorithm. Subsequently, we examined the correlation between the phylogenetic distance (i.e., whole genome substitution rate) of virus strains with spatial distance and the phylogenetic distance of their host individuals (measured as the substitution rate of the cox1 gene) using partial Mantel tests. Spearman correlation coefficients was employed in the partial Mantel tests. A significant positive correlation (p value < 0.05) indicates virus-host co-divergence or virus divergence through space.

## DATA AVAILABILITY

The assembled viral genome sequences have been deposited in the CNGBdb with the accession code N_AAACQU010000000-N_AAADML010000000 (see Supplemental Data 2). The sample metadata and other materials required to reproduce our computational and statistical results are provided in the GitHub repository along with code and scripts (XXXXXX).

## CODE AVAILABILITY

Code and scripts are provided in a GitHub repository (XXXXX).

## ACKNOWLEDGMENTS

This study was funded by grants from the National Key R&D Program of China (2021YFC2300900), National Natural Science Foundation of China (32270160), Shenzhen Science and Technology Program (JCYJ20210324124414040), and open project of BGI-Shenzhen Shenzhen 518000, China (BGIRSZ20210001). M. Shi was supported by Shenzhen Science and Technology Program (KQTD20200820145822023), Guangdong Province “Pearl River Talent Plan” Innovation, Entrepreneurship Team Project (2019ZT08Y464), and the Fund of Shenzhen Key Laboratory (ZDSYS20220606100803007). G. Liang was supported by the United States National Institutes of Health U01 (AI151810). We gratefully acknowledge colleagues at BGI-Shenzhen and China National Genebank (CNGB) for RNA extraction, library construction and sequencing.

## AUTHOR CONTRIBUTIONS

**Conceptualization**, DX Wang, JH Li, WC Wu and M Shi; **Methodology**, YF Pan, HL Zhao, QY Gou, DX Wang, JH Li, WC Wu and M Shi; **Sample Collection and Processing**, QY Gou, GY Luo, GY Xin, SJ Le, Jing Wang, X Hou, CH Yang, JX Cheng, YQ Liao, MW Peng, SQ Mei, JB Kong, Juan Wang, Chaolemen, YH Wu, JB Wang, TQ An, WC Wu; **Data analysis**, YF Pan, HL Zhao, QY Gou, PB Shi, DX Wang, JH Li, WC Wu and M Shi; **Writing – Original Draft**, YF Pan; **Writing – Review and Editing**, HL Zhao, QY Gou, ZR Ren, SQ Peng, JS Eden, J Li, B Li, DY Guo, GD Liang, X Jin, EC Holmes, DX Wang, JH Li, WC Wu, M Shi. **Funding Acquisition**, DX Wang, JH Li, WC Wu and M Shi; **Resources (sampling)**, JH Tian, Y Feng, K Li, WH Yang, D Wu, GP Tang, B Zhang, WC Wu; **Resources (computational)**, DX Wang, JH Li, M Shi; **Supervision**, DX Wang, JH Li, WC Wu and M Shi.

## COMPETING INTERESTS

The authors declare no competing interests.

## Notes

### Competing Interest Statement

The authors have declared no competing interest.

### Summary of Updates

uploading supplemental text and tables

