## Supplemental Information for "Metagenomic analysis of individual mosquitos reveals the ecology of insect viruses"

**Supplementary Figures**

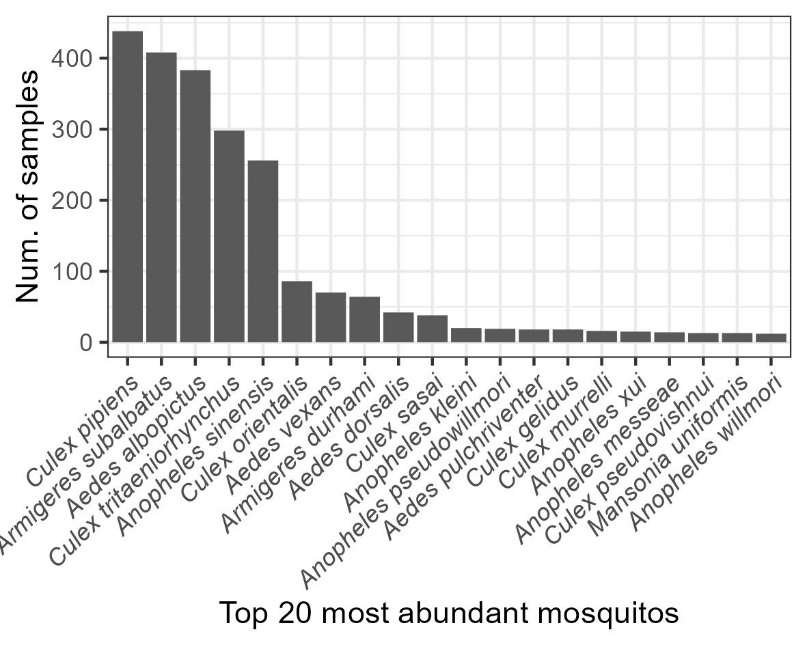

**Supplementary Fig. 1 | The abundance of the top 20 most abundant mosquito species.**

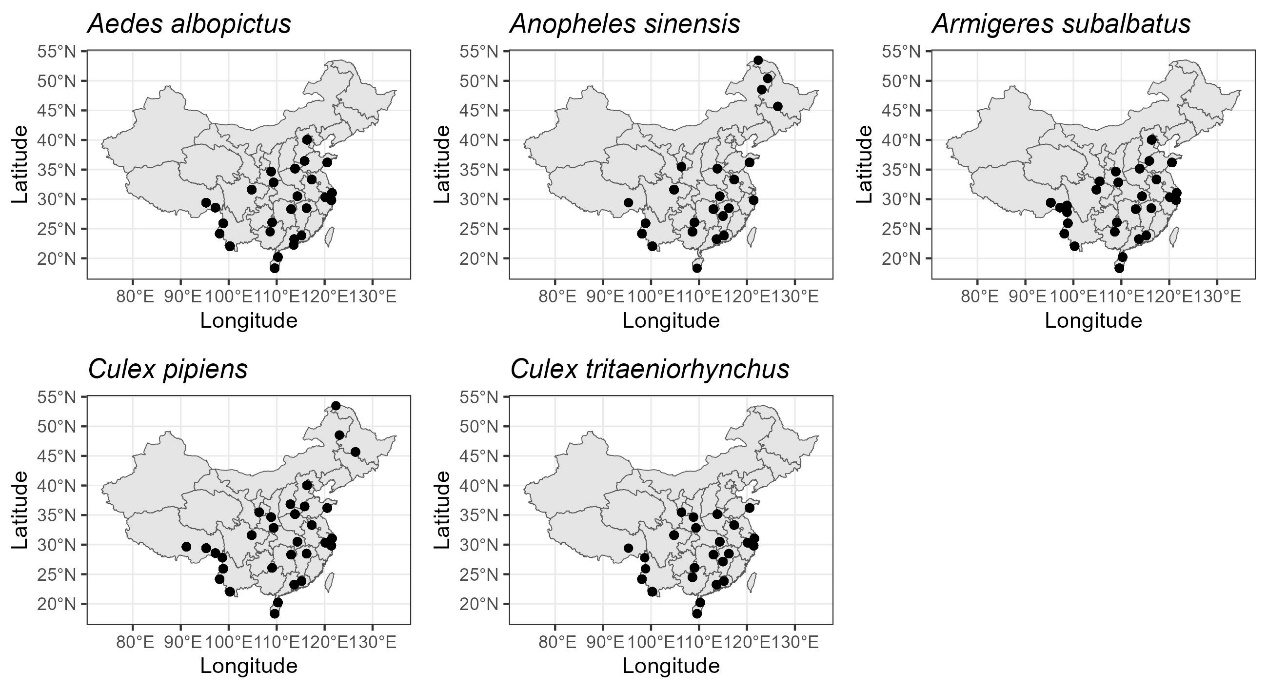

**Supplementary Fig. 2 | Distribution of the five dominant mosquito species in China.** Dots indicate incidence of the corresponding mosquito species.

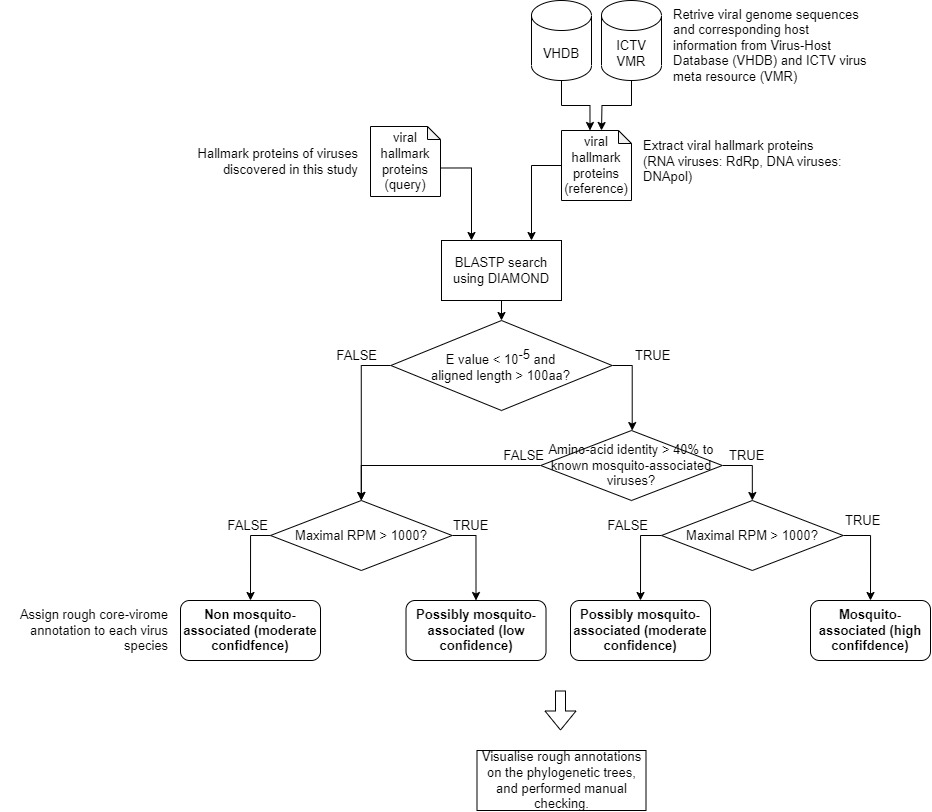
**Supplementary Fig. 3 | Flow chart of the core-virome identification process.**

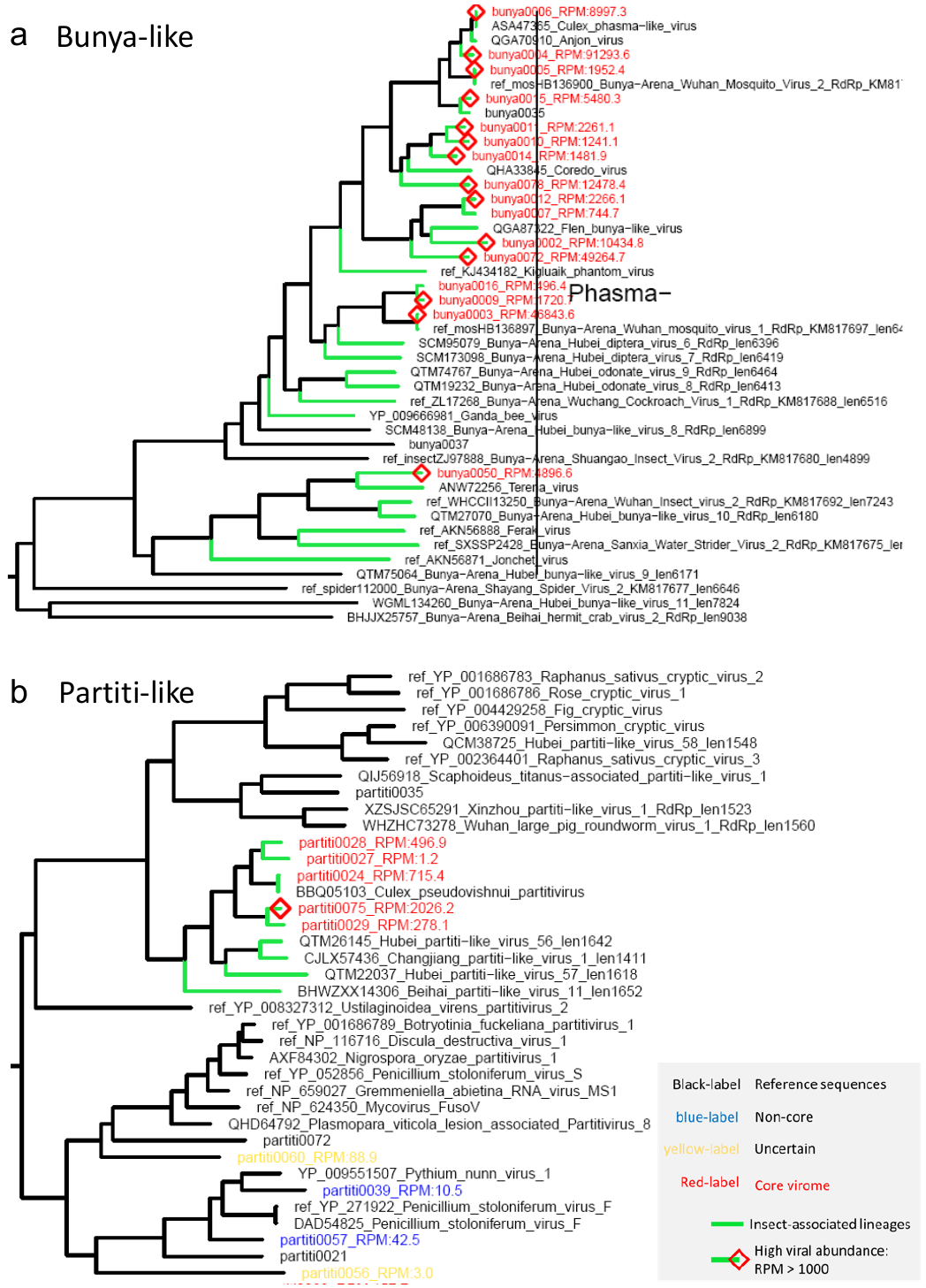

**Supplementary Fig. 4 | Examples of core-virome identification.** This figure illustrates the typical process to identify the core mosquito virome and phylogenetic trees for all viruses detected in this study is provided as Supplemental Data 3. **a,** the first example is a group of viruses from the bunya-like superclade. In the RdRp phylogeny we can confirm that core viruses belong to lineages known to be associated with insects, and the viral abundance were also very high. **b,** in the second example, we see that the red coloured viruses belong to mosquito associated lineage, so they were identified as core viruses. In contrast, the two blue coloured viruses belonged to fungi associated viral lineages, and we also detected high abundance of fungi in those samples, indicated that they are not core viruses. The remaining two yellow colour viruses are classified as uncertain, and were also excluded from downstream analysis. These uncertain viruses are too distantly related to any known viruses, and their abundance are low.

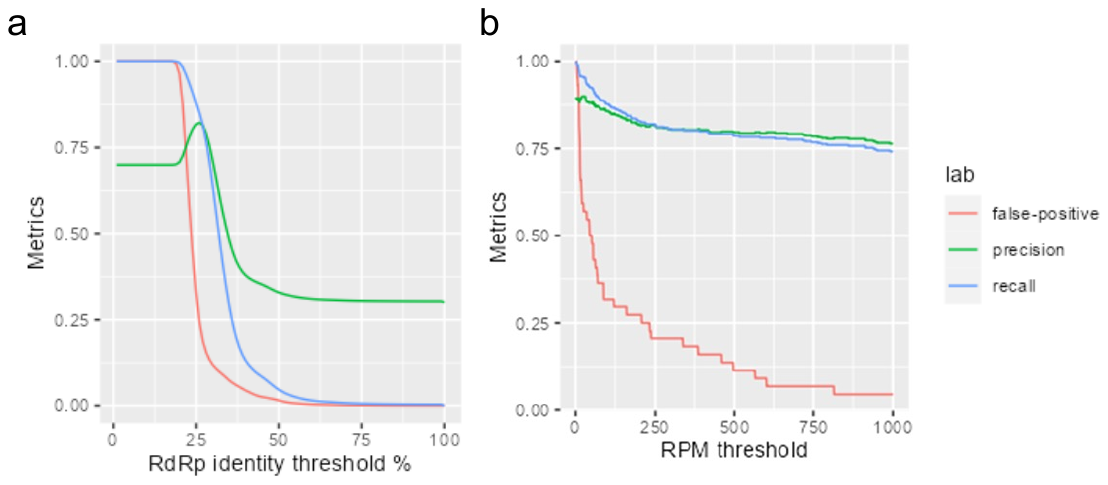

**Supplementary Fig. 5 | Benchmarking results for core-virome identification thresholds.** **a,** the benchmarking results for protein percentage identity threshold. **b,** the benchmarking results for viral abundance (measured by RPM) threshold.

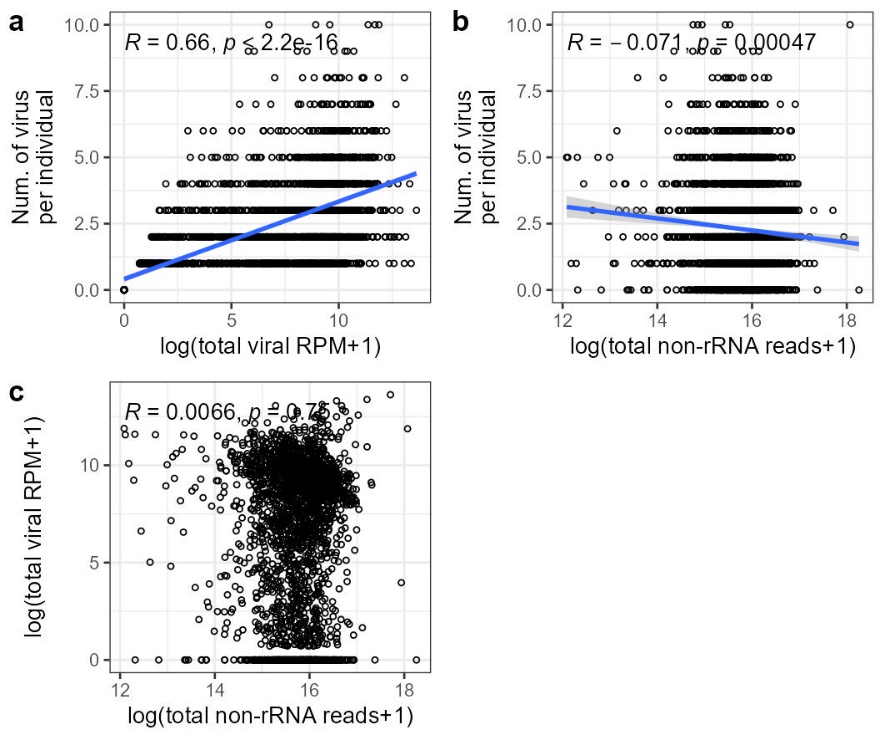

_­­­­­_**Supplementary Fig. 6 | The correlation between sequencing depth with viral abundance and richness. a,** the relationship between viral abundance and number of virus species per individual. **b,** the relationship between sequencing depth (measured as number of non-rRNA reads) and number of virus species discovered per individual. **c,** the relationship between sequencing depth and viral abundance. The statistics on the top of each plot show the Pearson correlation, where “*R”* is the correlation coefficient and “*p”* stand for p value. If there is significant correlation (p < 0.05), we visualized the trends with linear regression, which are shown as the blue lines, and shaded areas indicate 95% CI.

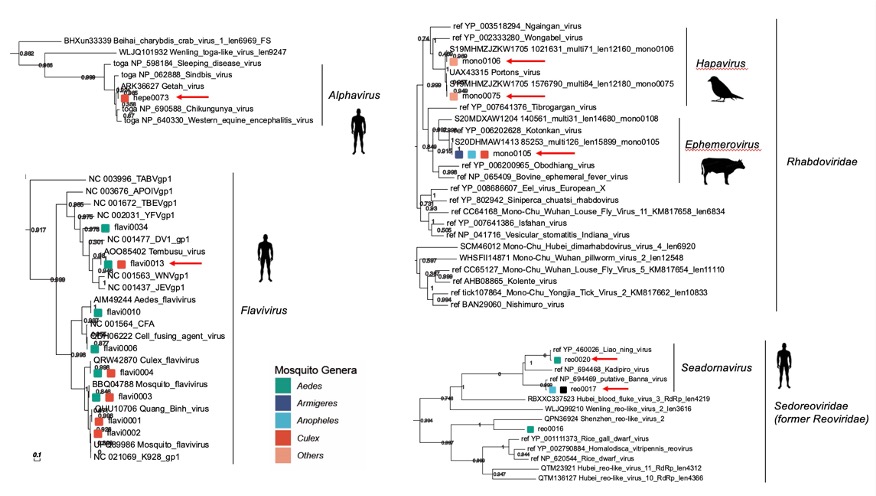

_­­­­­_**Supplementary Fig. 7 | Phylogenetic placement of the seven (putative) arboviruses.** The trees were estimated using protein sequences of RNA-directed RNA replicase, using maximal likelihood method. The seven arboviruses identified were marked with red arrows. The putative vertebrate hosts of corresponding viruses are visualised with cartoon.

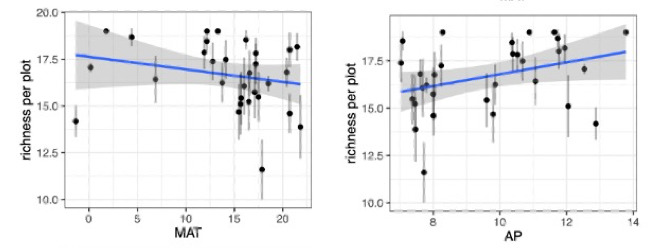

_­­­­­_**Supplementary Fig. 8 | Analysis of viral richness at mosquito population scale.** To robustly reflect the correlation between viral richness and environmental factors, we summarised the total viral richness of each mosquito population and analysed the correlation again. MAT: mean annual temperature, AP: annual precipitation.

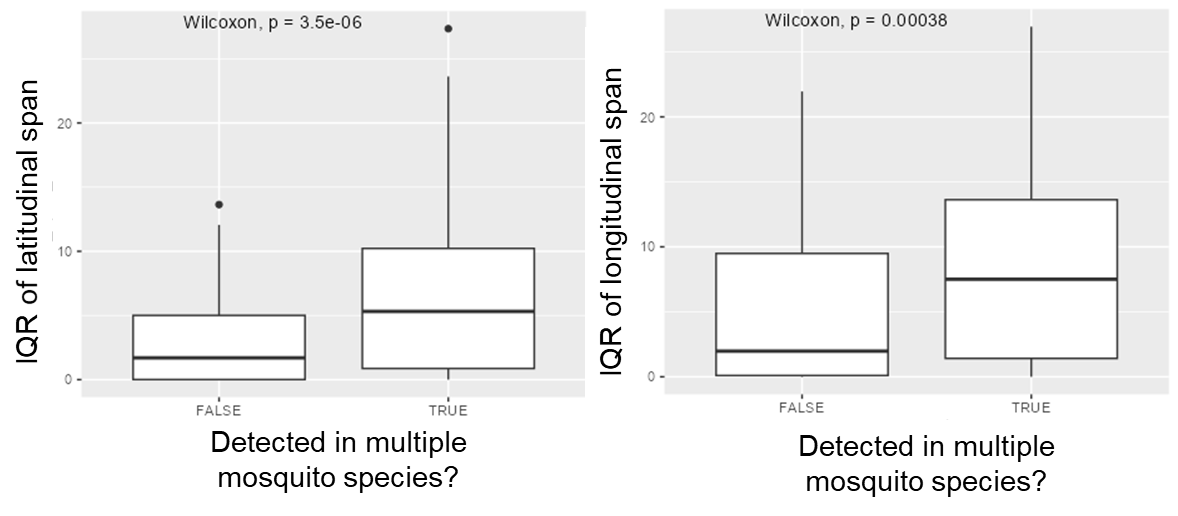

**Supplementary Fig. 9 | Comparison of distribution range between viruses shared among multiple mosquito species and viruses associated with a single species.**

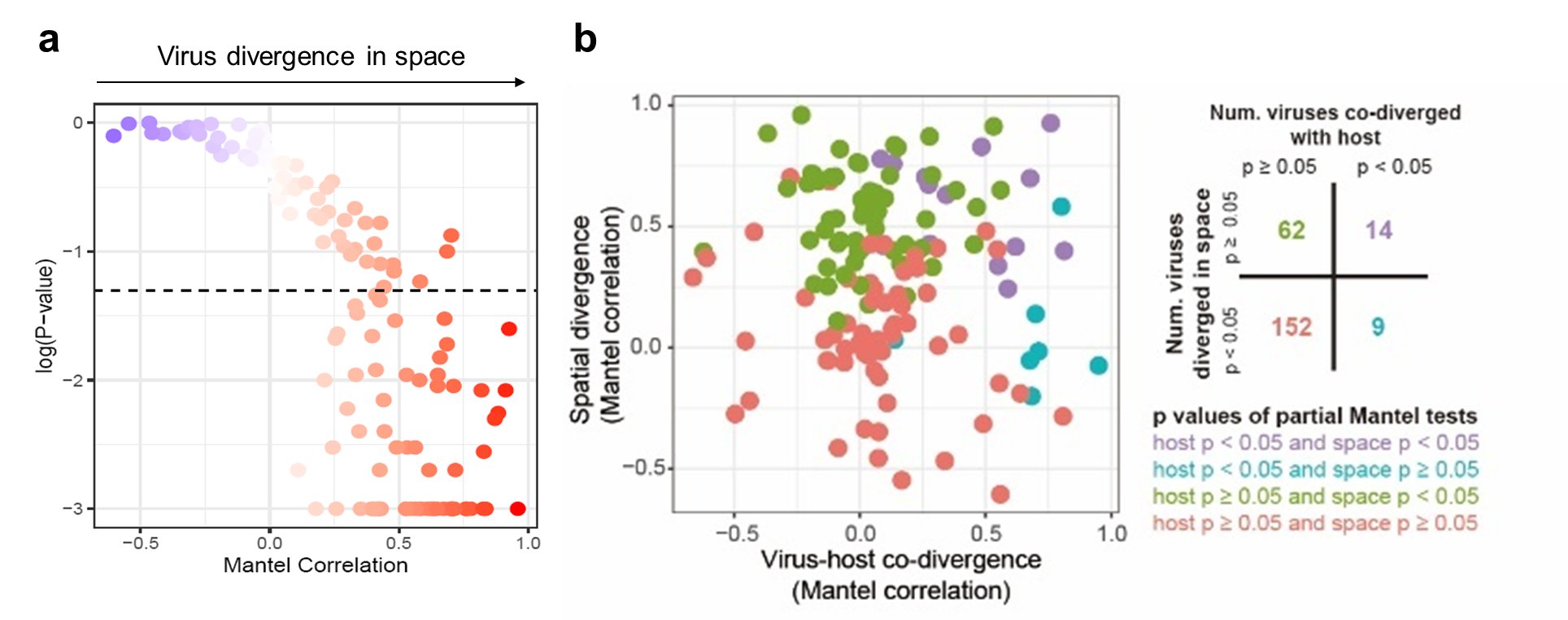

**Supplementary Fig. 10 | Genetic divergence of viruses in space. (a)** Trends of virus divergence in space. The degree of spatial divergence was measured using the Mantel correlation (Spearman) between the phylogenetic distance matrix of viruses and spatial distance. **(b)** The relation between trends of virus-host co-divergence and spatial divergence. A contingency table is shown on the right, where the numbers are virus species within each class.

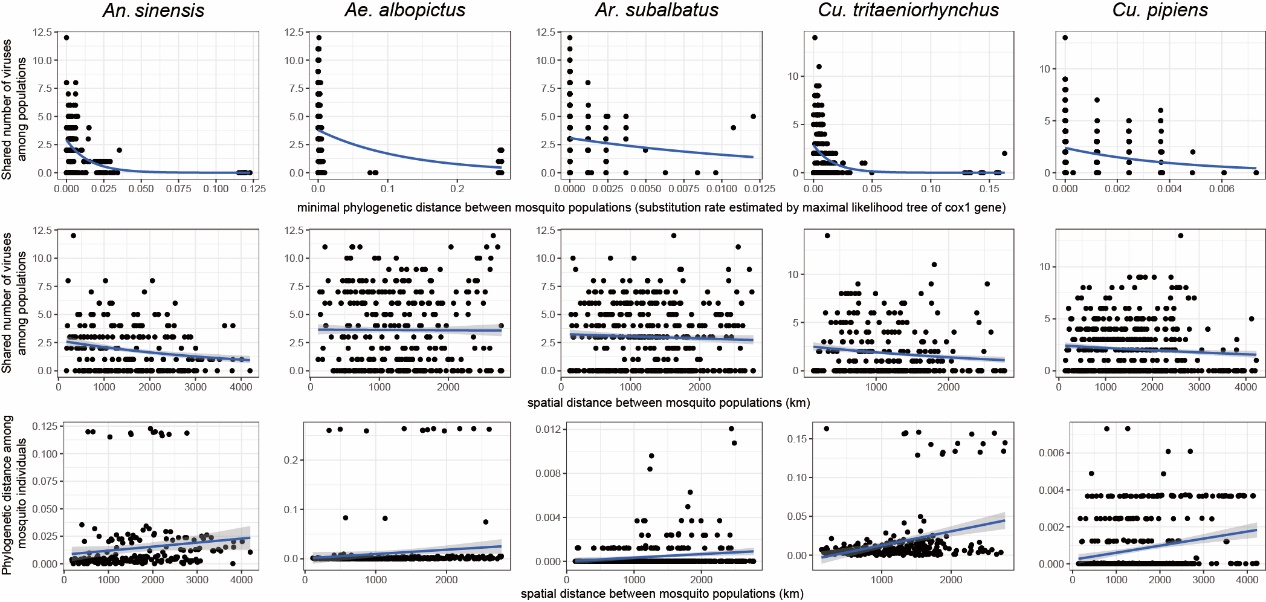

**Supplementary Fig. 11 | Comparison of distribution range between viruses shared among multiple mosquito species and viruses associated with a single species.**

**Supplementary Tables**

**Supplementary Table 1.** The distance-based redundancy analysis (dbRDA) models for virome composition at individual and mosquito population scale.

| **Individual or population** | **Distance Metrics** | **Best model structure supported by forward selection** | **Total variance explained by the best model** | **PERMANOVA p value for the best model** |
| --- | --- | --- | --- | --- |
| Individual mosquito | Jaccard | D ~ species + CPC1 + CPC2 + CPC3 + APC1 + APC2 + mammal_richness | 13.9% | <0.001 |
| ­­Individual mosquito | Bray-Curtis | D ~ species + CPC1 + CPC2 + CPC3 + APC1 + APC2 + mammal_richness | 17.8% | <0.001 |
| Mosquito population | Jaccard | D ~ species | 26.4% | <0.001 |
| Mosquito population | Bray-Curtis | D ~ species | 33.9% | <0.001 |

**Note:** “D” indicates dissimilarity matrix of viral community between mosquito individuals or populations. The variable “species” indicates mosquito species identity, “CPC1/2/3” stands for principal components of climate variables, “APC1/2” stands for principal components of arthome variables.

**Supplementary Table 2.** Contingency table of the number of virus species co-diverge with their hosts and spatially diverged with increasing spatial distance.

|  | **Not diverged through space** | **Diverged through space** | **Total** |
| --- | --- | --- | --- |
| **Not co-diverged with hosts** | 152 | 62 | 214 |
| **Co-diverged with hosts** | 9 | 14 | 23 |
| **total** | 161 | 76 | 237 |

**Supplementary Table 3.** Internal calibration points used to calibrate molecular clocks.

| **Calibration point** | **Node age** | **Reference** |
| --- | --- | --- |
| Diptera | 240-241 Mya | Krzeminski *et al*. 1994 |
| Culicidae | 90-99 Mya | Borkent & Grimaldi 2004 |
| Culicanae | 76.5-79 Mya | Poinar *et al*. 2000 |
